# Benchmarking of local ancestry inference with different assays and parameters

**DOI:** 10.64898/2026.05.18.726085

**Authors:** Tomoki Motegi, Franklin Huang, Joshua D. Campbell

## Abstract

Local ancestry inference (LAI) enables high-resolution characterization of chromosomal segments inherited from distinct ancestral populations, offering unique insights into genetic architecture in admixed cohorts. While LAI is commonly performed with high-coverage whole-genome sequencing (WGS), the ability of other genotyping assays or varying sequencing depths has not been thoroughly benchmarked. In this study, we systematically evaluated the accuracy of LAI across SNP microarrays, whole-exome sequencing (WES), and ultra low-pass WGS (ULP-WGS) using diverse validation samples and state-of-the-art imputation pipelines. We show that ULP-WGS, when paired with GLIMPSE2, achieves robust accuracy at 0.25x coverage with a minimum genome window size of 0.5 centimorgans, with mean accuracy minus one standard deviation exceeding 95%. For WES, using “on-target” reads alone yields suboptimal performance, particularly for European and South Asian ancestries with accuracy less than 79.1% and 70.6%, respectively. However, incorporating “off-target” reads in WES and utilizing GLIMPSE2 substantially improved accuracy ≥95% with a minimum window size of 0.2 centimorgans. We further evaluated formalin-fixed, paraffin-embedded (FFPE) samples and found that LAI could be performed successfully using WES data with accuracies of ≥95% at a minimum window size of 0.5 centimorgans. In contrast, SNP microarrays did not achieve substantial accuracies at any window size (≤95%). Together, these results demonstrate that LAI is achievable without conventional high-coverage WGS and establish optimal parameters for LAI across platforms.

## Introduction

Local ancestry inference (LAI), which estimates the ancestral origin of specific chromosomal segments in admixed individuals, has emerged as a powerful tool for investigating the genetic architecture of complex traits in diverse populations. Unlike global ancestry estimation that summarizes genome-wide admixture proportions, local ancestry (LA) provides high-resolution mapping of ancestry tracts, enabling the detection of ancestry-specific genetic effects that may be obscured in conventional genome-wide association studies (Suarez-Pajes et al. 2021). Recent advances in statistical and machine learning methods, including hidden Markov models and discriminative algorithms such as RFMix (Maples et al. 2013) and ELAI (Guan 2014), have substantially improved the accuracy and scalability of LAI using genotyping or sequencing data. Integrating LA into genetic analyses, such as admixture mapping, expression quantitative trait loci analysis, and heritability estimation, has been shown to enhance the detection of causal signals while mitigating confounding due to population structure (Zhong et al. 2019). Beyond methodological relevance, LA has enabled insights into population-specific disease mechanisms; for example, a study of Latin American lung cancer patients revealed that somatic *EGFR* mutations were more strongly associated with local Native American ancestry than with global ancestry, implicating ancestry-linked germline variation in the somatic evolution of cancer (Carrot-Zhang et al. 2021).

Despite these advances, a critical gap remains in the systematic evaluation of LAI across various genotyping platforms, including SNP arrays, whole-exome sequencing (WES), and whole-genome sequencing (WGS). Although it is well established that factors such as marker density, phasing accuracy, and reference panel composition influence LA accuracy (Suarez-Pajes et al. 2021), the ability to ascertain LA from different data modalities has not been rigorously benchmarked. Most existing studies have utilized high-density SNP arrays or WGS but have not assessed more cost-effective assays such as whole exome sequencing (WES) or ultra-low pass WGS (ULP-WGS) (Maples et al. 2013; Hilmarsson et al. 2021). Other factors can also influence the quality of genotype calls such as whether the DNA was derived from fresh frozen (FF) or formalin-fixed paraffin embedded (FFPE) tissues (Do and Dobrovic 2015; Guo et al. 2022). Overall, the lack of modality-aware benchmarking can hinder appropriate LAI on legacy datasets or in new sequencing projects that are utilizing low-cost assays.

In addition to data modality and reference resources, the accuracy of LAI is also strongly shaped by the choice of genomic window size. Most LA algorithms do not infer ancestry at individual variant sites. Instead, they aggregate information across contiguous genomic windows, which introduces an inherent trade-off between statistical stability and spatial resolution. Larger windows incorporate more SNPs, making ancestry estimates more robust and less sensitive to genotyping noise, phasing errors, or sparse marker distributions. However, large window sizes inevitably reduce resolution and may obscure short ancestry tracts or span recombination breakpoints. In contrast, smaller windows enable finer-scale delineation of ancestry transitions but rely on fewer informative markers and therefore incur greater variance and increased risk of misclassification. Critically, the optimal window size for LAI is not universal but is tightly coupled to the underlying assay and the number of genetic markers measures. This parameter has not been carefully examined when applying LA methods to heterogeneous or cost-constrained genomic datasets.

To address these limitations, our study systematically evaluated the performance of LAI algorithms across a diverse set of sequencing and genotyping platforms, including SNP arrays, WES, and ULP-WGS. By establishing best-practice guidelines tailored to each data type, we seek to enhance the accessibility, robustness, and generalizability of LA-based analyses in population and medical genomics, thereby enabling inclusive and biologically informed discovery across ancestrally diverse cohorts.

## Results

### Generation of synthetic datasets for LAI benchmarking

To perform benchmarking of key experimental factors that may influence LAI, we first needed to build a high-quality LA model and generate synthetic datasets representing different modalities with “ground truth” LA calls. A LA model was built with G-nomix (Hilmarsson et al. 2021) using a 30x coverage WGS dataset (Byrska-Bishop et al. 2022) consisting of 1,006 samples from four different global ancestral backgrounds taken from the 1000 Genomes project. LA calls were predicted on a validation set consisting of 62 samples which were used as the “ground truth” for all comparisons (**Figure 1A**). To create datasets for benchmarking, we generated simulated genotypes from the real WGS data in the validation set that mimic commonly used platforms, including ULP-WGS and SNP microarrays (**Methods**; **Figure 1B**). WGS data was down sampled to varying depths to create profiles like “low pass” and “ultra-low pass” (ULP) sequencing strategies. To generate microarray-like data, SNPs were limited to those specifically interrogated on the Affy6.0 SNP microarray (∼900K loci). To assess whole-exome sequencing (WES), we directly used the WES data provided by the 1000 Genomes Project on the same samples in the validation set. For datasets where read down-sampling occurred, *de novo* variant calling was performed. Each dataset then underwent phasing and imputation with Eagle-Beagle and/or GLIMPSE2 to generate final sets of genotypes for LA calling. LAI accuracy was quantified as the proportion of correctly inferred haplotype-level local ancestry calls across all genomic windows. Haplotype label ambiguity was accounting for by selecting the higher concordance between the two possible haplotype alignments. This benchmark design enabled us to systematically assess the influence of sequencing depth, SNP density, and modality on LAI accuracy.

**Figure 1.**
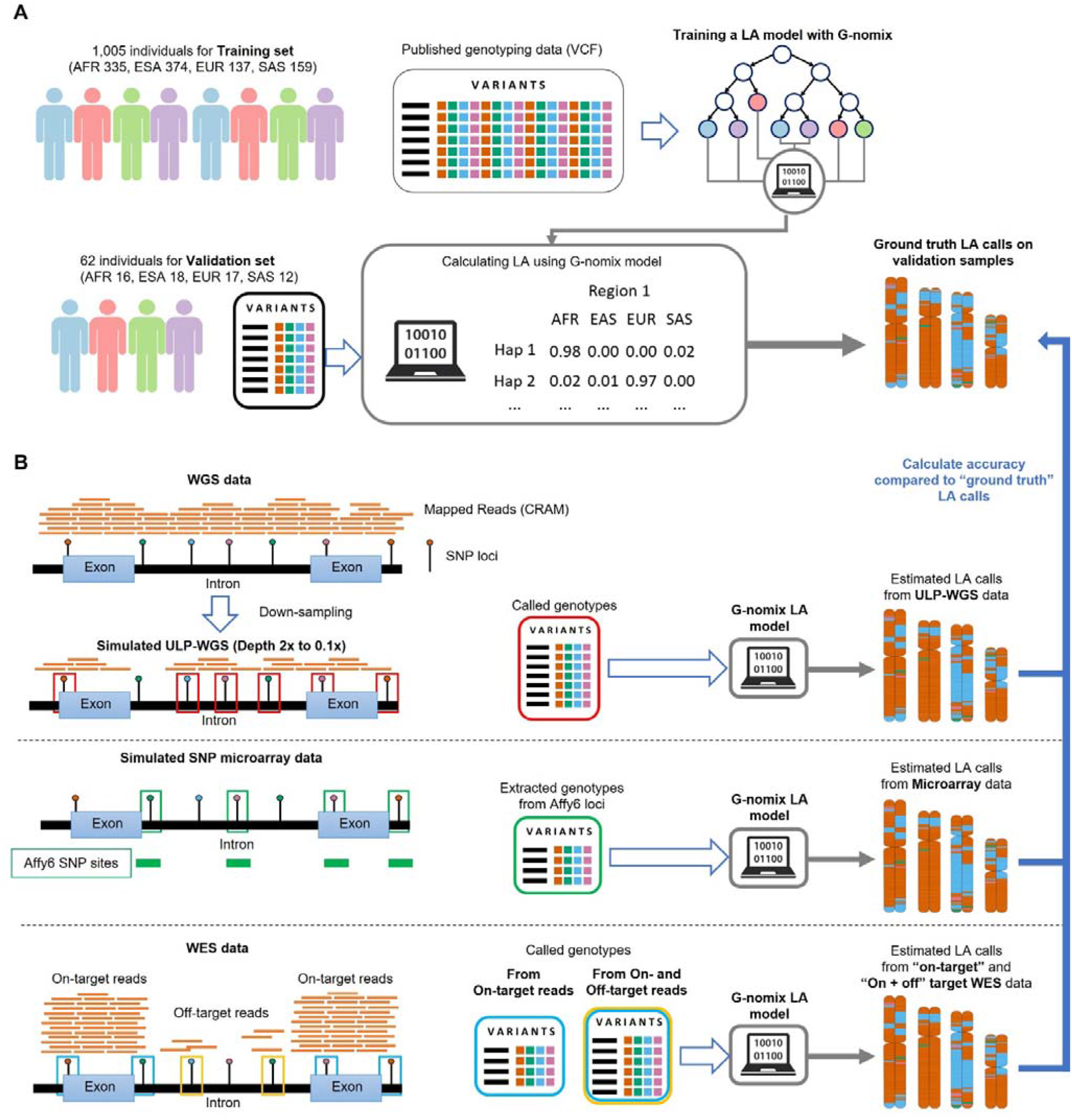
Generation of simulated datasets for benchmarking of local ancestry (LA) inference. **(A)** Generation of LA model and “ground truth” LA calls. 1,067 single-ancestry samples were selected from the 1000G project by ADMIXTURE clustering from published VCF data. 1,005 samples were assigned to the training set and the remaining 62 samples were assigned to the validation set. A distinct LA model was generated for each window size using the genotypes of the training set samples and G-nomix software. LA calls were generated for samples in the validation set using the G-nomix model and used as “ground truth” for all benchmarking comparisons. **(B)** Generation of simulated genotypes in validation samples to represent different modalities. Genotyping data was obtained or simulated to represent different sequencing modalities for samples in the validation set. Ultra-low pass WGS (ULP-WGS) data was simulated by down-sampling high-coverage WGS data. Microarray genotype data was simulated by limiting to the SNPs that matched the loci of the Affymetrix Genome-Wide Human SNP array 6.0 (Affy6). Real WES data was obtained for the samples in the validation set from the 1000G project. Variants from WES were called in using either “on-target” reads or both “on- and off-target” reads. Each set of variant calls underwent phasing and imputation with the Eagle-Beagle and/or GLIMPSE2 pipelines to obtain genome-wide genotypes. LA calls were predicted for the genotypes from each modality using the same G-nomix model generated in part A. Accuracy was determined for each modality by comparing the LA calls with those generated from full-coverage WGS data.

### Benchmarking LAI with ULP-WGS data

Utilizing our benchmarking framework, we examined varying sequencing depths and window sizes to ascertain how accurately LA calls can be recapitulated with low-pass and ULP-WGS data. Examples of the accuracy of LA calls at different sequencing depths are shown for chromosome 1 using GLIMPSE2-based imputation and a window size of 0.5 cM (**Figure 2A**). While the accuracy using ULP-WGS decreased with smaller window sizes and lower sequencing depths, a robust LAI performance (as a mean accuracy minus one standard deviation of ≥95%) was achieved at the smallest window size of 0.2 cM and a minimum sequencing depth of 0.25x (**Figure 2B**). At the lowest depth of 0.1x, a minimum window size of 8 cM was required to achieve an average of 95% accuracy although the variation in accuracy between samples was higher (**Figure 2B**). In contrast to GLIMPSE2, the popular Eagle–Beagle pipeline showed a pronounced decrease in performance under various sequencing depths. To achieve the robust accuracy threshold in ULP-WGS with Eagle phased-Beagle imputation, a minimum window size of 4 cM was required for a sequencing depth of 0.5x, a 2 cM window size was required for a depth of 1x, and a 1 cM window size was required for a depth of 2x (**Figure 2C**). The Eagle-Beagle pipelined failed to achieve average accuracies of 95% altogether at a depth of 0.1x.

**Figure 2.**
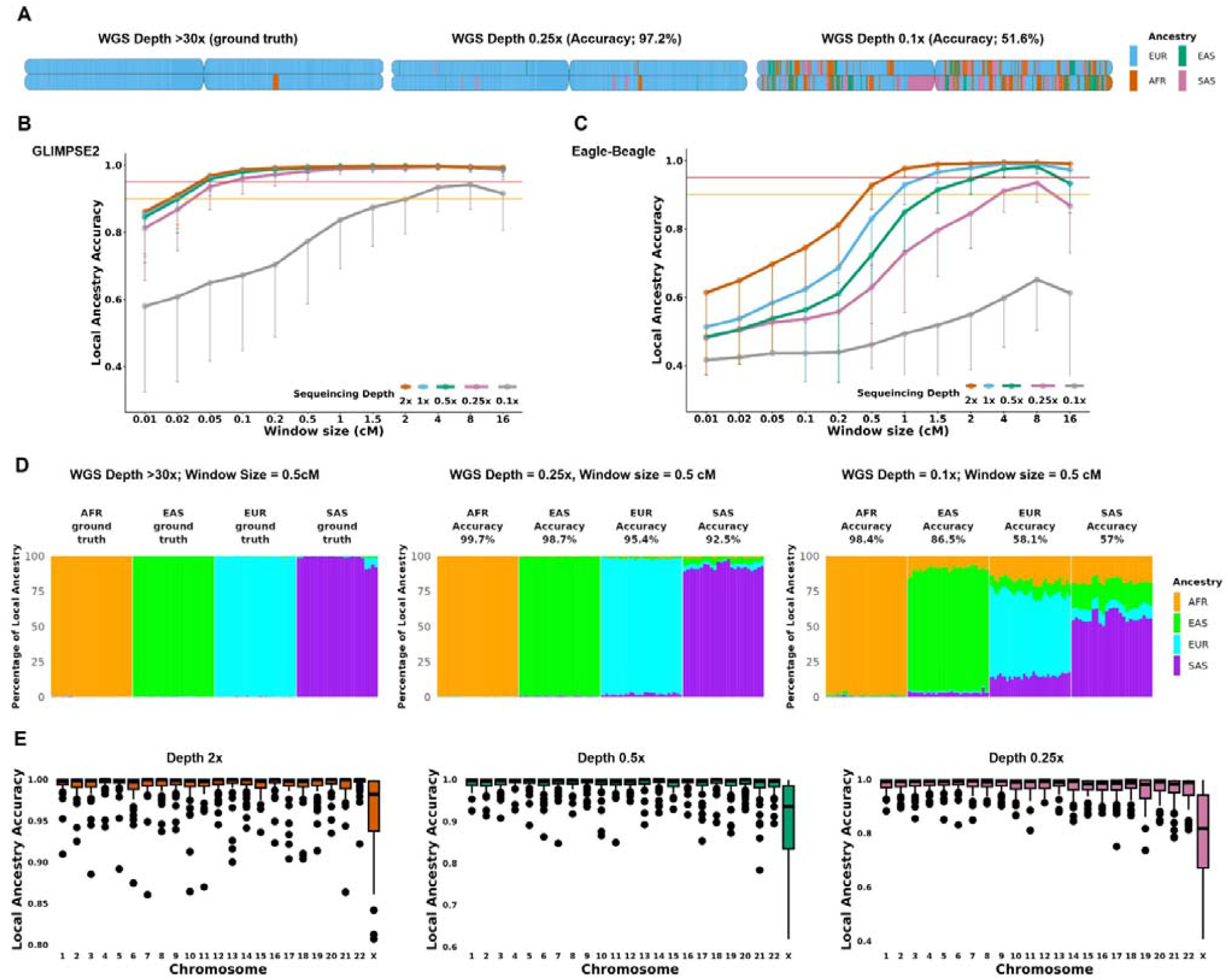
The effect of whole-genome sequencing depth on accuracy of local ancestry inference (LAI). **(A)** Examples of estimated ancestral blocks on chromosome 1 estimated by G-nomix in a single individual across different sequencing depths. True local ancestries were defined from published genotypes from 1000 Genomes Project and compared with results from genotypes called from various sequencing depths. Each block color indicated ancestry: blue for European, orange for African, green for East Asian, and purple for South Asian. **(B, C)** The accuracy of LAI by each window size on chromosome 1 using **(B)** GLIMPSE2 imputation and **(C)** conventional imputation, consisting of Eagle phasing followed by Beagle imputation. The y-axis represents the average accuracy with LAI as determined by G-nomix from the established genotypes in the 1000 Genomes Project, while the x-axis shows the window sizes in centimorgans (cM). The horizontal red line demarcates an LAI accuracy of 0.95, whereas the orange line demarcates an LAI accuracy of 0.90. The vertical line descending from each LAI point indicates the range to the lower limit of standard deviations (SD). **(D)** The percentage of local ancestral haplotypes on chromosome 1 are shown for each sample in the validation dataset within each sequencing depth. Results are shown for 0.5 cM windows. **(E)** Chromosome-level LAI accuracy for individual samples are shown for sequencing depths of 0.25x, 0.5x, and 2x at a window size of 0.5 cM . Each box represents the distribution of LAI accuracy across individual samples for each autosomal chromosome, with the center line indicating the median, and the box denoting the interquartile range. Individual points beyond the whiskers were plotted as outliers.

Notably, the accuracy varied by ancestry, with African ancestry demonstrating the highest estimation accuracy, followed by East Asian, European, and South Asian ancestries. At 0.25× sequencing depth with a 0.5 cM window, the mean accuracies were 99.7%, 98.7%, 95.4%, and 92.5% for African, East Asian, European, and South Asian ancestry, respectively. This trend was observed consistently across sequencing depths (**Figure 2D**). Finally, we examined the accuracy of LA calls for different sequencing depths across chromosomes to determine if particular regions are more difficult to characterize. Chromosome-wise local ancestry accuracy revealed robust performance on the autosomes, whereas chromosome X exhibited lower estimation accuracy (**Figure 2E**). Specifically, GLIMPSE2-based imputation preserved exceptionally high accuracy (>98%) even at sequencing depths as low as 0.25× and with fine-scale window sizes (0.2–0.5 cM) for the autosomes. These findings indicate that ULP-WGS, when paired with modern imputation methods such as GLIMPSE2, can deliver reliable local ancestry inference at lower sequencing depths, supporting its utility as a cost-effective strategy for LA analyses in large-scale genomic studies.

### Performance of LAI in SNP Microarray and WES

For SNP microarray (Affy6) and on-target WES data, we employed the conventional Eagle-Beagle pipeline for imputation. Although we also tested GLIMPSE2 for these datasets, the mean LAI accuracy did not exceed approximately 0.27 for either Affy6 or on-target-only WES. This poor performance was likely due to GLIMPSE2 being optimized for low-coverage whole-genome sequencing data with continuous genome-wide read information, whereas Affy6 and on-target-only WES are marker-based assays with sparse or discontinuous genomic coverage. Also, this distinction was particularly important for on-target-only WES, where marker SNPs are confined to captured regions and are more sparsely and unevenly distributed than in SNP microarrays. Despite yielding similar numbers of imputed variants, LAI based on Eagle–Beagle imputation showed lower accuracy and greater variability for on-target WES compared with Affy6 at the same window size (**Table 1**). Specifically, while Affy6 genotypes on chromosome 1 achieved a robust LAI performance for window sizes larger than 0.5 cM (**Figure 3A**), on-target-only WES failed to reach this threshold at any window size.

**Table 1.** The parameters of Imputation for WES and Affy6.

|  | Affy6 | WES On-target | WES Joint |
| --- | --- | --- | --- |
| Pre-phasing | Eagle | Eagle | GLIMPSE2 |
| Imputation | Beagle 5.4 | Beagle 5.4 | GLIMPSE2 |
| Number of Imputed Variants | 1,970,122 | 1,970,122 | 3,751,808 |
| Number of variants with Impute2-like INFO scores >0.3 | 1,715,089 | 1,491,086 | 3,751,808 |
| Number of intersecting variants | 557,752 | 555,299 | 558,956 |
| Model variants covered | 99.78% | 99.35% | 100.00% |
| Window size | 2.0 cM | 2.0 cM | 2.0 cM |
| Mean Accuracy | 0.994 | 0.892 | 0.983 |
| SD | 0.011 | 0.088 | 0.032 |

**Figure 3.**
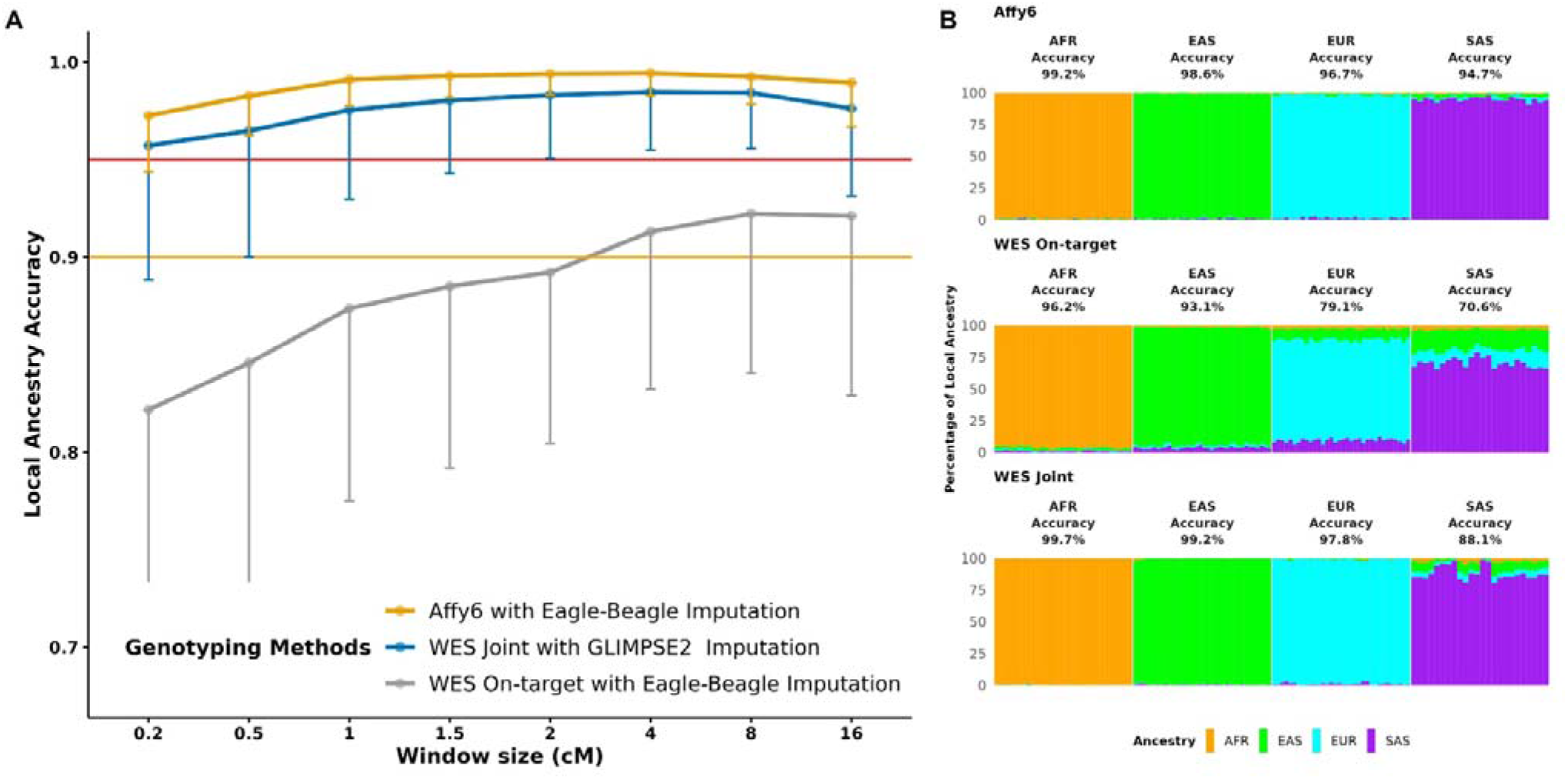
The accuracy of LAI by diverse genotyping methods on chromosome 1. **(A)** The LAI accuracies through sparse genotyping modalities are shown for the Affy6 paired with Eagle for phasing followed by Beagle for imputation (yellow line), whole exome sequencing (WES) on-target paired with Eagle-Beagle imputation (gray line), and WES on-target and off-target (Joint) with GLIMPSE2 imputation (Blue line). The y-axis represents the average accuracy with LAI as determined by G-nomix from the established genotypes in the 1000 Genomes Project, while the x-axis shows the window sizes in centimorgans (cM). The horizontal red line demarcates an LAI accuracy of 0.95, whereas the orange line demarcates an LAI accuracy of 0.90. The vertical line descending from each LAI point indicates the range to the lower limit of standard deviations (SD). **(B)** The percentage of local ancestral haplotypes on chromosome 1 are shown for each sample in the validation dataset within each genotyping modality. Results are shown for 2 cM windows.

Most WES protocols rely on hybrid capture to select DNA from regions of interest (e.g. exons). However, these hybrid capture methods are often imperfect and reads from “off-target” regions are captured, albeit to lower extents. As off-target reads are randomly distributed across the genome, they mimic coverage observed in ULP-WGS. For example, the average coverage of off-target reads in WES data in the 1000G validation sample set was 1.16x. Indeed, incorporating off-target regions into WES and using GLIMPSE2 for imputation substantially improved LAI performance at a window size of 2 cM (**Figure 3A**). This joint approach yielded over twice as many high-quality imputed variants compared to the on-target method alone, elevating the mean accuracy from 0.892 to 0.983, and reduced the standard deviation, thereby demonstrating robust LAI performance with lower variability across samples (**Table 1**). Consequently, strategically repurposing off-target reads transforms standard WES from a suboptimal assay for LAI into a highly accurate, standalone modality.

To determine if these improvements applied universally, we compared the accuracy of ancestry-specific estimation across the platforms. Affy6 data yielded stable estimates closely matching the ground truth, with only a slight reduction for South Asian subjects (**Figure 3B**). On-target-only WES produced reliable estimates for African and East Asian ancestries but showed consistently reduced performance for European and South Asian groups. At a 0.5 cM window, the mean accuracies were 96.2%, 93.1%, 79.1%, and 70.6% for African, East Asian, European, and South Asian ancestry, respectively. However, the inclusion of both on-target and off-target reads in WES markedly enhanced accuracy in these groups, resulting in estimates comparable to those from Affy6 for African, East Asian, and European ancestries. By elevating WES accuracy to match Affy6 across African, East Asian, and European ancestries, this joint approach successfully mitigates platform-induced biases, ensuring more equitable and robust LAI in diverse cohorts.

### Impact of formalin-fixed, paraffin-embedded (FFPE) Preservation on Ancestry Estimation Accuracy

FFPE samples represent one of the most abundant resources for genomic studies of retrospectively collected clinical samples, particularly in oncology, as they enable the long-term preservation of clinical material. However, fixation and paraffin embedding are known to fragment and chemically modify DNA, leading to reduced read coverage and genotyping call rates and lower reliability of downstream analyses. These artifacts raise the concern that LAI from FFPE-derived DNA may be less accurate compared to fresh-frozen or blood-derived samples, especially when using array-based genotyping platforms. Given the widespread reliance on FFPE tissue in research, it is important to clarify how preservation affects ancestry estimation. To this end, we analyzed four African and four European samples that were each profiled with four distinct modalities from the TCGA prostate adenocarcinoma (PRAD) dataset (Affy6-Fresh-Frozen, Affy6-FFPE, WES-Blood, and WES-FFPE). The data from the Affy6 Fresh Frozen assay was used to generate the “ground truth” local ancestries for all comparisons. We observed that FFPE samples processed with Affy6 exhibited markedly reduced accuracy in LA calls, never achieving greater than 95% accuracy at any window size. In contrast, WES FFPE samples showed high concordance with their blood-derived counterparts (**Figure 4A**). An ancestry-specific breakdown further revealed that FFPE preservation introduced notable inaccuracies in both African and European ancestry estimates in Affy6 data. In contrast, WES-based estimates remained stable and consistent across ancestral groups (**Figure 4B**). Together, these results underscore the platform-dependent sensitivity of LAI to fixation artifacts and indicate that sequencing-based approaches provide a more robust strategy for recovering accurate ancestry information from archival FFPE tissues.

**Figure 4.**
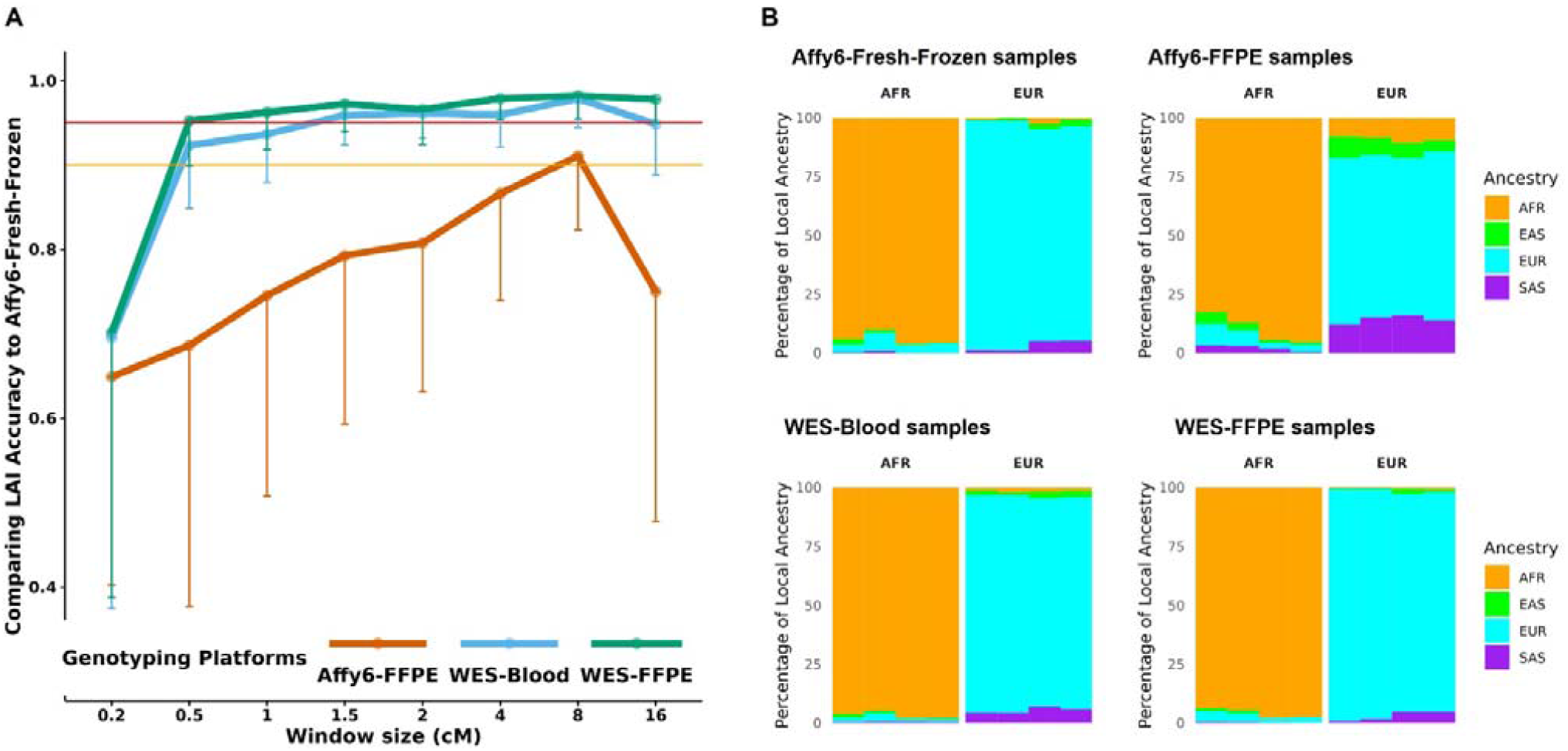
Impact of Formalin-Fixed, Paraffin-Embedded (FFPE) fixation on LA accuracy. **(A)** LAI accuracy on chromosome 1 across sample types and genotyping platforms was shown for FFPE samples analyzed with Affy6 (Affy6-FFPE; orange line), non-fixed blood samples with WES (WES-Blood; blue line), and FFPE samples with WES (WES-FFPE; green line). The y-axis indicates concordance with LAI estimates derived from Affy6-based fresh frozen samples (Affy6-Fresh-Frozen) using G-nomix, while the x-axis represents window sizes (centimorgans; cM). The red and orange horizontal lines indicated LAI accuracy thresholds of 0.95 and 0.90, respectively. Vertical lines from each data point represented the lower bound of the standard deviation (SD). **(B)** Haplotype-level LAI accuracy on chromosome 1 was evaluated across sample types (non-fixed and FFPE) and genotyping methods (Affy6 and WES) using four TCGA samples. Stacked bar plots depicted LAI results within 0.5 cM windows, including two individuals of African ancestry (AFR) and two of European ancestry (EUR).

## Discussion

LAI is increasingly recognized as a powerful approach for investigating ancestry-specific genetic architecture in admixed populations, with applications ranging from admixture mapping to ancestry-aware eQTL and GWAS analyses (Suarez-Pajes et al. 2021). However, despite its growing importance, most benchmarking studies have focused on high-coverage WGS, with limited attention to more cost-effective or legacy genotyping platforms such as SNP microarrays, WES, or ULP-WGS. In this study, we systematically evaluated the performance of LAI across these diverse modalities using ancestry-stratified validation datasets and a range of analytical settings. Notably, our results show that ULP-WGS, when combined with modern imputation tools, can achieve LAI accuracy comparable to that of standard WGS. Specifically, GLIMPSE2-based imputation maintained high accuracy even under very low sequencing depths and small window sizes, whereas the conventional Eagle–Beagle pipeline showed substantially reduced performance under the same conditions. These findings are consistent with prior studies, which demonstrate that imputation under sparse data conditions, such as ULP-WGS or ancient DNA, is inherently challenging due to elevated genotype uncertainty, reduced phasing accuracy, and dependence on the composition of the reference panel (Hui et al. 2020). Furthermore, recent benchmarking has demonstrated the superior performance of GLIMPSE2, particularly in recovering variants and maintaining accuracy at a biobank scale (Rubinacci et al. 2023). Collectively, our results underscore that while LAI from ULP-WGS is feasible, its success relies heavily on the use of imputation frameworks specifically optimized for low depth of coverage data. When paired with carefully selected tools and parameters, ULP-WGS offers a scalable and cost-effective strategy for analysis of local ancestry, particularly in large and ethnically diverse cohorts.

We also found substantial differences in LAI performance across genotyping platforms. Pseudo-microarray data based on SNPs from the Affy6 array achieved high accuracy (>95%) across most ancestries, though slight reductions were noted for South Asian individuals. WES data limited to on-target regions performed reasonably for African and East Asian ancestries but showed reduced accuracy for European and South Asian groups. Prior studies have shown that combining WES with SNP microarrays and imputation can yield high concordance in global ancestry inference and capture rare coding variants with power comparable to whole-genome sequencing (WGS), as demonstrated in the UK Biobank study (Barton et al. 2021). However, these hybrid approaches have limitations for high-resolution LA estimation. In addition to their reliance on multiple data types, which increases analytical complexity and cost, such designs are not compatible with legacy datasets or biobanked samples for which only WES or targeted panel sequencing data are available. As a result, while they may improve global ancestry inference, their practicality and scalability for retrospective studies and diverse cohorts remain limited. Building on the approach of a previous study, which enhanced LAI by incorporating off-target reads in exome sequencing (Hu et al. 2013), we treated off-target WES regions as analogous to ULP-WGS, increasing marker density genome-wide. Coupled with GLIMPSE2-based imputation, this strategy significantly improved LAI accuracy in WES data, exceeding 95% at a 2□cM window. Notably, unlike many existing approaches that rely on large population-scale datasets to compensate for sparse genomic coverage, our benchmarking demonstrates that accurate local ancestry can be inferred from WES assays alone, without the need for joint array data or biobank-level sample sizes. Together, these findings demonstrate that despite the limitations of WES, for local ancestry, extending WES with off-target data and optimized imputation pipelines can yield accurate inference of local ancestries in retrospective tissue biobanks.

An additional and practically relevant aspect of our study was the evaluation of FFPE samples, which are commonly used in retrospective genomic research, particularly in oncology. Consistent with previous reports demonstrating reduced genotyping call rates and accuracy in FFPE samples processed via SNP arrays (Thompson et al. 2005; Hosein et al. 2013), we observed a marked reduction in LAI accuracy with FFPE-derived array data. In contrast, WES-derived FFPE samples showed high concordance with matched blood-derived WES data, indicating that sequencing-based approaches are more robust to FFPE-induced DNA degradation. These findings highlight the platform-dependent sensitivity of LAI to fixation artifacts and underscore the importance of selecting the appropriate assay when working with archival tissue. Importantly, our results demonstrate that accurate LAI is achievable from FFPE samples when using optimized sequencing workflows, supporting the feasibility of analysis of LA in cancer research where FFPE tissues are common. The ability to recover reliable LA information from FFPE-derived WES data will enable the study of ancestry-associated tumor biology and contribute to more inclusive frameworks for precision oncology.

This study has three potential limitations. First, although we used high-quality public datasets, such as the 1000 Genomes Project and TCGA, these may not fully capture the technical variability and population diversity encountered in real-world settings. However, their consistency and standardization provide a reliable framework for benchmarking. Second, we focused exclusively on G-nomix for LAI. While it demonstrated strong performance, future comparisons with other algorithms such as RFMix (Maples et al. 2013) and ELAI (Guan 2014) may elucidate further improvements to the accuracy of LAI in specific settings. Lastly, our FFPE analysis was limited to four prostate cancer samples. Additional cohorts that have run genomic assays on matched FFPE tissue and blood will be needed to more broadly support the feasibility of LAI in archival tissues in other settings.

Taken together, this work provides a modality-aware framework for selecting and optimizing LAI strategies. We demonstrate that the combination of data type, sequencing depth, window size, and imputation method has a significant impact on LAI accuracy. Our benchmarking offers practical guidance for researchers working with diverse sample types and resource settings, thereby enabling robust and equitable integration of LAI into genomic studies of complex traits and diseases.

## Methods

### Data acquisition and sample selection

We selected 1,067 single-ancestry samples from the 1000 Genomes Project, as defined by unsupervised ADMIXTURE clustering following previously reported criteria (Hilmarsson et al. 2021), from different global ancestral backgrounds. 1,005 samples were stratified into a training set (335 African Ancestry; AFR, 374 East Asian Ancestry; EAS, 137 European Ancestry; EUR, and 159 South Asian Ancestry; SAS) while 62 samples were used for the validation set (16 AFR, 18 EAS, 17 EUR, and 12 SAS). The 1000G Phase 3 genotype dataset in VCF format was used for training, whereas whole-genome sequencing (WGS) and whole-exome sequencing (WES) CRAM files were employed for the validation dataset (http://ftp.1000genomes.ebi.ac.uk/). To assess the effect of LAI on Formalin-fixed, Paraffin-embedded (FFPE) samples, we selected four patients from The Cancer Genome Atlas Prostate Adenocarcinoma (TCGA-PRAD) dataset who had both fresh-frozen and FFPE samples derived from the same specimen. WES BAM files and Affymetrix Genome-Wide Human SNP Array 6.0 (Affy6) binary files in their native Affymetrix formats were obtained from the NCI’s Genomic Data Commons (GDC) portal (https://portal.gdc.cancer.gov/).

### Estimation of “ground-truth” local ancestries from 1000G genotype data

The overview of the methodology is in Figure 1A. A high-quality LA model was built using the full coverage WGS data. LAI was conducted using G-nomix, which combines methodologies from historically successful LAI classifiers, including Random Forest and Support Vector Machines and has demonstrated improved accuracy in ancestry calling compared to RFMix (Maples et al. 2013). We constructed a LAI model using training data from the published 1000G VCFs using the default mode in G-nomix (Logistic regression + XGBoost). Subsequently, this model was applied to predict the LA of the samples in the validation set. These inferred local ancestries on the validation set were used as the reference (i.e. “ground truth”) for all benchmarking comparisons. G-nomix was run using different window sizes from 0.01 to 16 centimorgans (by modifying the smooth_size parameter) to obtain ground truth LA calls at different resolutions.

### Simulation of ULP-WGS genotypes

We performed down-sampling of high-coverage WGS data to simulate low pass (i.e. 1x) or ultra-low pass (i.e. 0.1x) coverage and assess how different depths of coverage affect the accuracy of LAI. The original WGS CRAM files were down-sampled to sequencing depths of 0.1x, 0.25x, 0.5x, 1x, and 2x using the subsample option in Samtools view (version 1.12). Variant calling was performed to retain all SNPs with genotype likelihoods using optimized Bcftools (version 1.12) mpileup (-q 20 -Q 20 -I -E) and Bcftools call (-Aimv -C alleles) commands.

### Simulation of SNP microarray genotypes

Affy6 genotype data was simulated for the validation samples by extracting SNPs corresponding to the 903,073 loci in the SNP 6.0 annotation file that realigned to GRCh38 using the published VCF dataset (Figure 1B). FFPE genotypes were determined using the birdseed-v2 algorithm (Korn et al. 2008), and the resultant birdseed files were converted into VCF format using the +affy2vcf (The gtc2vcf software tool. https://github.com/freeseek/gtc2vcf. Accessed Jan 17 2023.) in bcftools plugins with the realigned annotation file.

### WES genotypes

We incorporated two types of genotype data: standard genotypes calling from WES on-target sites and expanding genotypes amalgamating on-target and off-target reads to increase the accuracy of the WES data. On-target regions were defined using the capture BED annotation for Illumina’s Rapid Capture Exome kit v1.1, as provided by the Broad Institute, which specifies the genomic intervals targeted during exome enrichment. Genotypes for the on-target WES dataset were generated exclusively from variants falling within these capture regions. To generate the expanded WES dataset, both on-target and off-target reads were retained and processed jointly using the same method as in the Simulation of ULP-WGS genotypes section. In this framework, off-target reads arising naturally from hybrid capture produced substantially lower and more uneven coverage (average 1.16x) than on-target regions (average 62x).

### Genotype imputation

We conducted genotype imputation to improve LA estimation and integrate genotypic data across different platforms. Using the training dataset, we constructed a reference panel for imputation and haplotype phasing. Quality control was performed using BCFtools, which involved removing rare variants including singletons and doubletons, splitting multiallelic sites, aligning variants to the reference genome, and removing duplicate and missing genotypes. Haplotype phasing was carried out using a reference-based approach with Eagle (version 2.4) with optimized parameters (--Kpbwt 20000). Post-phasing, chromosome-wise genotype imputation was conducted using Beagle (version 5.4) with a reference panel derived from the 1000 Genomes Project Phase 3 dataset and converted to bref3 format. Imputation parameters were evaluated with reference to previously reported (Pook et al. 2020), and the following parameters were used: ne = 20, window = 80, overlap = 8, and iterations = 30. Imputation tools like Beagle fail to produce definitive genotype calls in low-coverage sequencing, as they do not explicitly model genotype likelihoods derived from sparse sequencing reads. We also employed GLIMPSE2 (Rubinacci et al. 2023), a recently developed imputation software tailored for ULP-WGS data imputation. The GLIMPSE2 pipeline consisted of the following steps using default parameters. First, the “GLIMPSE2_chunk” defined imputation regions across entire chromosomes. Next, “GLIMPSE2_split_reference” generated binary reference panels from quality-controlled reference data. Subsequently, the “GLIMPSE2_phase” was used to conduct genotype imputation across genomic segments using default parameters. Finally, the “GLIMPSE2_ligate” merged the imputed genomic chunks by each chromosome.

### Evaluation of accuracy for LA calls

Following genotype imputation, LA was predicted using the G-nomix models for each window size built on the training dataset. The smooth size was adjusted to match the LAI window size because larger genomic windows yield fewer total windows, and excessively large smoothing parameters can exceed the available number of windows during model training. Accordingly, we used smooth sizes of 50 for window sizes up to 0.5 cM, 30 for 1.0 cM, 15 for 1.5 and 2.0 cM, 8 for 4.0 cM, 4 for 8.0 cM, and 2 for 16.0 cM. Individual LA predictions were compared against the “ground truth” LA calls for each haplotype within each chromosomal window. For each sample, accuracy was defined as the proportion of windows in which the inferred ancestry label matched the true ancestry label, averaged across the two haplotypes. Because haplotype phase labels (paternal vs. maternal) may be arbitrarily assigned, accuracy was computed under both possible haplotype correspondences between inferred and true LA calls, and the higher of the two concordance values was retained. For example, with a window size of 1 cM, chromosome 1 was partitioned into 286 windows, resulting in 286 windows × 2 haplotypes per window, or 572 total LA calls per sample. Sample-level accuracy was therefore defined as the proportion of correct LA calls out of 572. The mean and standard deviation were calculated to assess accuracy across different genotyping methods, and to account for sample-to-sample variability, we further applied a conservative criterion in this study, defining robust performance as conditions in which the mean accuracy minus one standard deviation was ≥95%. Global ancestry was determined as the modal (most frequent) LA state across all windows and chromosomes for each individual.

## Data Access

The 1000 Genomes Project Phase 3 data used in this study were obtained from the official FTP repository (http://ftp.1000genomes.ebi.ac.uk/vol1/ftp/phase3/). Whole-exome sequencing (WES) and SNP microarray data for TCGA prostate adenocarcinoma (TCGA-PRAD) samples were accessed via the Genomic Data Commons (GDC) Data Portal (https://portal.gdc.cancer.gov/) following dbGaP authorization. All data were processed by the relevant data use agreements and institutional guidelines.

## Funding

This work was conducted as part of the National Cancer Institute (NCI) Center to Reduce Cancer Health Disparities (CRCHD) R01CA248920 to J.D.C. and F.H.

## Competing Interest Statement

None declared.

## Ethics approval

Ethical approval is not applicable for this article.

## Acknowledgments

We are grateful to members of our laboratory and collaborators for helpful discussions and technical support. Computational resources were provided by BU Shared Computing Cluster. TM was supported by the Japan Society for the Promotion of Science, KAKENHI Grant Numbers 25K23673 and 26K18196.

## Contributors

Study Design: T.M., J.D.C

Data Analysis: T.M.

Manuscript drafting: T.M., J.D.C, F.H.

## Notes

### Competing Interest Statement

The authors have declared no competing interest.

